# Genotyping-by-sequencing and DNA array for genomic prediction in soybean oil composition

**DOI:** 10.1101/2024.06.07.598034

**Authors:** Melina Prado, Regina Helena Geribello Priolli, Evellyn Giselly De Oliveira Couto, Felipe Sabadin, Kaio Olimpio das Graças Dias, José Baldin Pinheiro

## Abstract

Soybean oil is intended for various purposes, such as cooking oil and biodiesel. The oil composition changes the shelf life, palatability, and how healthy this oil is for the human diet. Genomic selection jointly uses these traits, phenotypes, and markers from one of the available genotyping platforms to increase genetic gain over time. This study aims to evaluate the impact of different genotyping platforms, DNA arrays, and Genotyping-by-Sequencing (GBS) on genomic selection in relation to the composition of fatty acids in soybean oil and total oil content. We used different quality control parameters, such as heterozygote rate, minor allele frequency, and missing data rate in ten combinations, and two prediction models, BayesB and BRR. To compare the impact of the genotyping approaches, we calculated the principal components analysis from the kinship matrices, the SNP density, and the traits prediction accuracies for each approach. Principal component analysis showed that the DNA array explained better the population genetic architecture.

On the other hand, prediction accuracies varied between the different genotyping platforms and only GBS was affected under different quality control parameters. Although the DNA array has important and well-studied polymorphisms for soybeans and is stable, it also has ascertainment bias. GBS, although not stable and requires more robust quality control, can discover alleles specific to the population under study. As soybean oil is used for different functions and the fatty acid profiles are different for each objective, the work constitutes a critical study and direction for improving the composition of soybean oil.

## Introduction

One of the main questions in genomics is understanding how genetic variability influences the range of different natural traits. There are many ways to genotype individuals. However, the most common way in modern genetics is using abundant Single Nucleotide Polymorphisms (SNPs), such as Genotyping-By-Sequencing (GBS) and DNA array. SNP genotyping platforms made it possible to establish genomic selection as a routine method in breeding programs, in which a dense and wide distribution of markers is necessary for good accuracy (MEUWISSEN; HAYES; GODDARD, 2001). In this context, the DNA array genotyping platform enables the extraction of individual genomic information with greater ease, precision, and speed, showing differences in translocations, inversions, and polymorphisms in just one nucleotide (SONG et al., 2020).

DNA arrays, addressing allelic polymorphisms, are tailored to specific species, which means that many organisms, mainly native and orphan crops, still lack arrays accessible on the market. The narrower the genetic base of an organism, the more likely a high-efficiency array of genotyping in the populations that will be used for selection. The other side is that, as they represent the most representative polymorphisms of a given species, many rare SNPs of great importance for a specific population are not captured with a DNA array, and this is information to be taken into account when choosing the genotyping approach (Albrechtsen et al., 2010). For species that have a broad genetic base and consequently ascertainment bias by the use of SNP arrays or even the lack of a published reference genome, an exciting way of genotyping is Genotyping-By-Sequencing (GBS) (ELBASYONI et al., 2018).

GBS is a technique that discovers markers at the same stage as the genotyping (CROSSA et al., 2013). The technique is simple, has an excellent cost-benefit ratio compared to other genotyping methods, and produces high accuracies when used in genomic prediction (JARQUíN et al., 2014). The technique can identify SNPs and *InDels* that may not be recognized on a DNA array, as the discovery of the markers is performed for the population under study, not in a general way as it is in commercial DNA arrays. It can also be applied to organisms with a reference genome and those without (ELSHIRE et al., 2011; SABADIN et al. 2022). As it has a wide range of possible outcomes, confidence is also lower compared to well-established arrays on the market. More robust bioinformatics analysis is needed since the approach may have a lot of missing data and erroneous SNPs (GLAUBITZ et al., 2014). To address these problems, quality control (QC) and imputation of SNPs are necessary.

Raw data after genotyping are generally unreliable, so quality control methods are effective for controlling Type I and Type II errors, false positives, and false negatives, respectively. This control happens with the removal of SNPs suspected of being errors or even with removing a sample that was not well genotyped and had a high rate of missing SNPs. Typically, the first step is filtering by call rate of genotypes, then filtering by MAF (*Minor Allele Frequency*), followed by filtering by missing data. Applying filters such as heterozygote rate and deviation from the Hardy-Weinberg equilibrium is still possible. While there are common pipelines for performing quality control, each filtering parameter is specific to the study population. The filters change according to the type of the population, whether they are self-pollinating or open-pollinating crops, the type of genotyping, whether by DNA array or Genotyping-By-Sequencing (GBS) approach, the size of the population, with the quality of genotyping, among others (PAVAN et al., 2020). Even in genomic selection in soybeans, there are different filters and genotyping approaches in the published works (for more details, see Supplementary Material 1).

Genomic selection can accelerate the soybean breeding process by reducing the time of its cycles through the early selection of genotypes, even before their phenotypes are evaluated. However, several factors influence the predictive accuracies, such as the markers’ density, the training sets’ size and composition, statistical prediction models, and the genetic architecture of the target trait (CROSSA et al., 2017; ISIDRO et al., 2015). Studies with genomic selection in soybeans are not as frequent as in maize. One of the first studies was from Jarquín et al. (2014), resulting in an accuracy of 64% for soybean yield. Most studies on genomic selection in soybeans focus on total oil, protein content, and some essential fatty acids. Still, to our knowledge, no study has been carried out simultaneously with genomic prediction for the five principal fatty acids.

Soybeans are one of the world’s main oilseeds, and of all the products made from soybean seeds, oil is the main one. Soybean oil represents 27% of the total oil produced worldwide and 27% of the total oil used for domestic consumption (USDA, 2022/23). Soybean oil is composed of five principal fatty acids, which are stored in the cytosolic lipid droplets of the seeds; they are palmitic acid (16:0), stearic acid (18:0), oleic acid (18:1), linoleic acid (18:2) and linolenic acid (18:3). The work of Yao et al. (2020) showed that both the total content of soybean oil and the percentage of fatty acids are quantitative traits with locus of major effect. The content of these acids gives soybean oil different characteristics according to the needs of each end user; it is possible to obtain a soybean oil that is healthier for blood health, has composition characteristics for use as biodiesel, or has a longer shelf life or even one that has superior performance for baking applications (FEHR, 2007). The difficulty in improving oil composition lies in the complexity of the fatty acid metabolic pathway. In addition to the fact that all fatty acids are correlated and belong to the same metabolic route (SONG; TAYLOR; ZHANG, 2023), they impact many agronomic characteristics of soybeans, mainly on mechanisms related to stress conditions, development, and plant growth (BELLALOUI; REDDY; MENGISTU, 2015; ZHAO; KOSMA; LÜ, 2021).

Although there are numerous possibilities for improving the composition of soybean oil, some studies are needed aimed at a better balance between fatty acids so that there are no significant impacts on the crop’s agronomic traits. Increasing oleic acid can decrease grain size and lower the germination rate while increasing saturated acid content can result in dwarf plants and abnormal leaf morphology. Increasing linolenic acid can reduce tolerance to abiotic stresses, and decreasing linolenic acid can lead to a decrease in total oil content, among many other problems associated with changing soybean oil composition. (SONG; TAYLOR; ZHANG, 2023).

This work aims to compare the impact of two genotyping approaches, SNPs from GBS and DNA arrays, in different predictive models of genomic selection for soybean oil composition under various quality control parameters.

## Material and methods

### Plant Material and field experiments

To represent the diversity of soybean oil composition, 94 common accessions were used (PRIOLLI et al., 2019). Of these 94 accessions coming from germplasm from Embrapa/Soja and the Department of Genetics (ESALQ/USP), 62 were Plant Introduction (PI), coming from China, Korea, Japan, and India, and 32 were unrelated cultivars (C) coming from Brazil and the United States of America. The seeds were arranged in an augmented block design (Federer, 1956) between November and March in two consecutive years, in the years 2009-2010 and 2010-2011, with each year containing six blocks and each block containing a maximum of 20 treatments. Each treatment consisted of 20 plants arranged 0.8 meters apart.

### Phenotypic traits

We extract the total soybean oil from each sample’s macerated bulk of 100 grams of seeds. We used the hexane solvent by the Butt method at the Instituto Agronômico de Campinas (IAC) in São Paulo, Brazil. Fatty acid profiles were constructed using gas chromatography (Chromatograph model 3900, Varian, Palo Alto, C.A), and Pearson’s correlation was calculated for all the traits. The six traits evaluated in the work were total oil, oleic acid (18:1), linoleic acid (18:2), linolenic acid (18:3), stearic acid (18:0) and palmitic acid (16:0) (PRIOLLI et al., 2019).

### Genotyping methods

We used data from two genotyping approaches, BarcSoySNP6k genotyping (Illumina Innum BeadChips) (SONG et al., 2020), hereafter referred to as “SNP_6K”. In addition to the one generated by genotyping-by-sequencing, hereafter referred to as “SNP_GBS”. For both methods, accessions were planted in germination trays, and the DNA was extracted from a five plant leaf bulk from each accession using the DNeasy Plant Kit (Qiagen). The trays were placed in a greenhouse at 24-25 degrees Celsius and 33% humidity. Both genotyping was performed at the ESALQ/USP Functional Genomics Center in Piracicaba, São Paulo, Brazil.

The GBS genotyping method was performed using two types of restriction enzymes, one with rare cut, the *Nsi*I enzyme (New England BioLabs Inc. ®), and another with frequent cut, the *Mse*I enzyme (New England BioLabs Inc. ®). Single-end sequencing was performed at the Blood Center of Ribeirão Preto, São Paulo, Brazil, using the Illumina NextSeq 500 platform (Illumina, San Diego, CA, USA). The sequencing quality of the two approaches was verified using a diagnostic tool of the generated data, the FastQC software v. 0.11.5 (Brabahan Bioinformatics). For the SNP calling, we used the TASSEL-GBS pipeline (GLAUBITZ et al., 2014) using *Willians 82v*.*4* as a reference genome and *Bowtie2* software (LANGMEAD; SALZBERG, 2012). To assess the difference in the distribution of the markers across the soybean genome in the two genotyping approaches, we plotted the marker’s density per base pair using the *mRVB* package (YIN et al., 2021). Finally, to observe the effect of different genotyping approaches on population structure, we constructed a kinship heatmap using the *AGHmatrix* package (AMADEU et al., 2023) and a Principal Component Analysis graph.

### Quality control parameters and imputation

In order to understand the predictive accuracy behavior under different combinations of quality control parameters, we used three distinct filters: I) Minor Allele Frequency (MAF) with maximum values of 1 and 5%; II) Missing Data (MD) with maximum values of 10, 20, 30 and 45%; and III) Heterozygotes rate (Het) with maximum values of 10 and 20%. To make the study discussion more straightforward, the combination of filters used will be cited, as shown in Table 1.

**Table 1.**
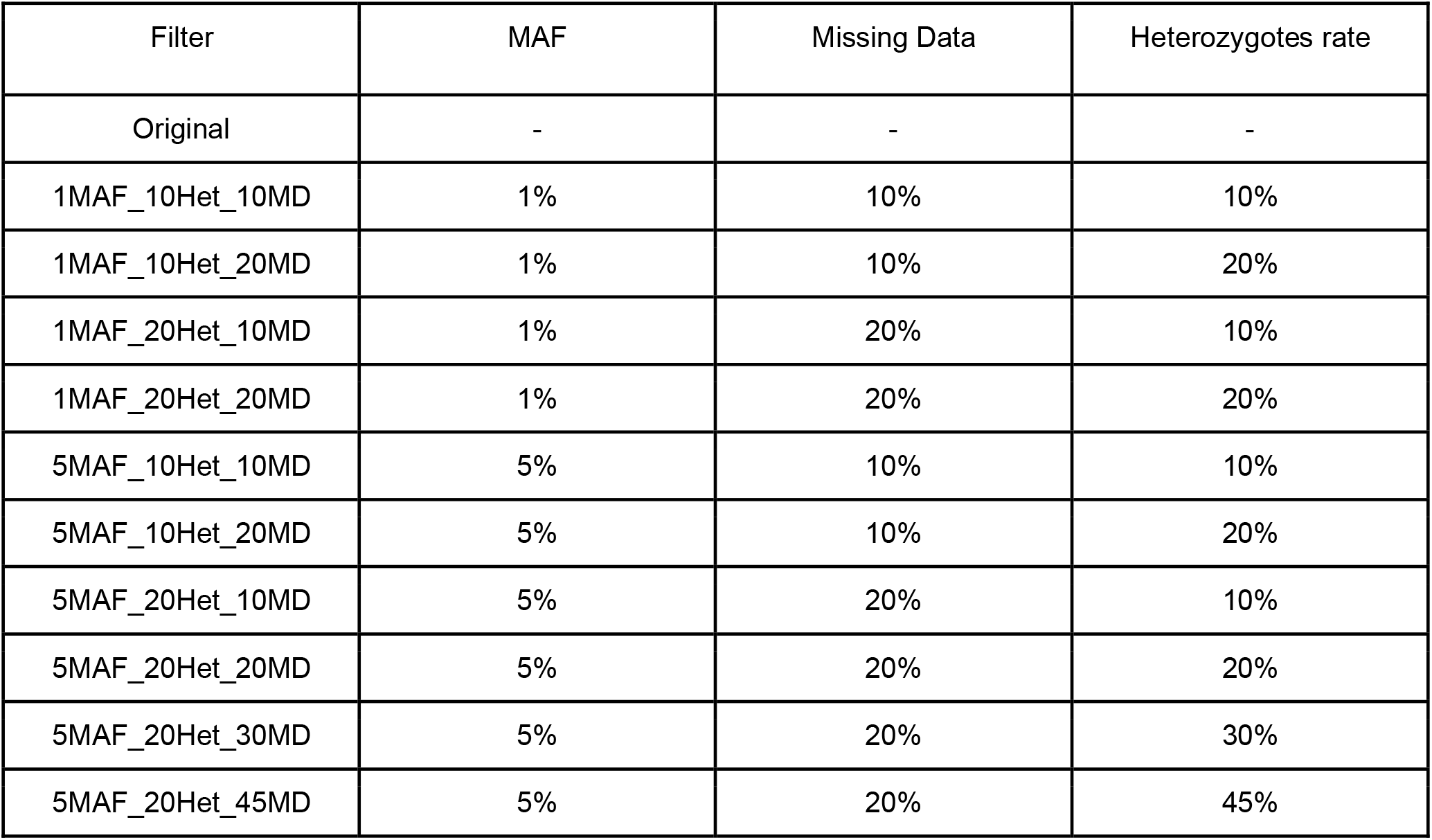
Different quality control filters are used to study genotype data processing in soybean oil traits in two genotype platforms: DNA array and GBS.

After the filtering step, in both genotyping approaches, SNP imputation was performed to prepare the data for genomic prediction analyses using *Beagle v. 5*.*1* (BROWNING; BROWNING, 2007).

### Single-trait genomic prediction

The genotypes Best Linear Unbiased Estimators (BLUEs) were estimated using the REML/BLUP method (*Residual Maximum Likelihood/Best Linear Unbiased Predictor*) using the *AsReml* package in the R environment (version 4.0.1, https://www.r-project.org/). The first model used was:

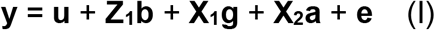

Where **y** is the vector that represents the average phenotype at the field plot level; **b** is the vector of random block effect, where 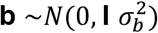; **g** is the vector of genotype fixed effect; **a** is the vector of year fixed effect; **e** is the vector of residual random effect, where 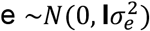; **X**_**1**_ and **X**_**2**_ are the incidence matrices for **g** and **a**, respectively; **Z**_**1**_ is the incidence matrix for **b** effects, and **I** is an identity matrix.

To estimate the variance components and the generalized heritabilities (Cullis et al. 2006), in addition to the genotypes BLUPs, the second model was used:

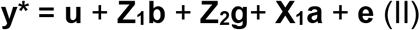

In model (II), **y\*** is the adjusted means vector or BLUEs calculated in model (I); all the other effects were estimated as was stated in model (I), except for the effect of genotype “**g**”, which was considered random, where 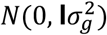. The Likelihood Ratio Test (LRT) was applied to test the genetic variance components.

The genomic selection models used were the BayesB and Bayesian Ridge Regression (BRR) methods. Both models are Bayesian approaches, but BayesB assumes that there may be many loci with no effect on the trait considered and that the loci that affect the trait follow a negative chi-square distribution (MEUWISSEN; HAYES; GODDARD, 2001). Meanwhile, the BRR method performs a homogeneous shrinkage between all markers and has a Gaussian prior density (DE LOS CAMPOS et al., 2013). Both models used the “BGLR” package in the R environment. In addition to performing a cross-validation step (10 k-fold), the analyses were repeated 10 times (PÉREZ; DE LOS CAMPOS, 2014). Predictive accuracy (PA) was calculated as the correlation between predicted genomic values and adjusted means. The BayesB model used 10000 iterations and 5000 BurnIn as parameters.

## Results

### Statistical analysis in the phenotypic data

Statistical analysis revealed that all genomic components calculated in the oil traits were statistically significant by the LRT (Table 2). The adjusted means and standard deviations showed variation within and between the oil traits. In this studied soybean population, the most abundant fatty acid was linoleic acid, followed by oleic, palmitic, linolenic, and stearic acids. The genetic variance values varied from 0.35 in stearic acid to 57.50 in oleic. The residual variance was higher in the oleic acid (15.50) and lower in the stearic acid (0.09). The coefficient of variation varied from 2.88 (stearic acid) to 37.20 (oleic acid). Considering all the oil traits, the oleic and linoleic acids had the highest genetic and residual variance and the highest coefficients of variation. Regarding Cullis heritability, values varied from 0.74 in stearic acid to 0.95 in palmitic acid, showing great potential for obtaining high accuracies in trait prediction. Furthermore, the heritability values of the traits, in decreasing value order, were for palmitic acid, followed by total oil, linoleic, oleic, linolenic, and stearic acid.

**Table 2.** Statistical analysis of 94 soybean oil content and composition accessions was evaluated for two years under Brazilian field conditions. Check’s and Regular’s average BLUEs represent the value of mean BLUES. Values in parentheses represent the standard deviation. Parameters such as Genetic Variance (GV), Residual Variance (RV), the Coefficient of Variation (CV), the Heritability of Cullis (Hc), and values for the Likelihood Ratio Test (LRT) were calculated for each trait.

| Trait | Check's average BLUES | Regular's average BLUES | GV | EV | CV | Hc | LRT |
| --- | --- | --- | --- | --- | --- | --- | --- |
| Linoleic | 53.10<br>(49.3 - 55.7) | 52.80<br>(28.20 - 61.11) | 35.00 | 9.02 | 28.40 | 0.89 | 121* |
| Linolenic | 6.38<br>(5.19 - 7.54) | 6.36<br>(2.41 - 11.59) | 1.77 | 0.70 | 7.95 | 0.83 | 76* |
| Oil | 21.00<br>(20.5 - 21) | 18.80<br>(12.90 - 23.30) | 5.24 | 0.93 | 5.13 | 0.91 | 133* |
| Oleic | 23.80<br>(19.90 - 30.10) | 24.60<br>(13.30 - 56.20) | 57.50 | 15.50 | 37.20 | 0.88 | 117* |
| Palmitic | 11.00<br>(10.50 - 11.50) | 10.60<br>(3.51 - 15.51) | 2.67 | 0.26 | 4.89 | 0.95 | 218* |
| Stearic | 3.00<br>(2.48 - 3.41) | 3.22<br>(2.16 - 4.55) | 0.13 | 0.09 | 2.88 | 0.74 | 44.6* |

Notes:
\* Significance when $p < 0.01$

Pearson’s correlation, distribution, and phenotypic classes of the analyzed oil traits can be found in **Figure 1**. The highest positive correlation observed was between linoleic and linolenic acids, with a value of 0.57. Conversely, the highest negative correlation observed was between oleic and linoleic acids, with a value of -0.94. Palmitic acid statistically correlated with all traits except for total oil (-0.074). Meanwhile, stearic acid showed a significant correlation only with palmitic acid (0.11) and total oil content (-0.11), although the correlations were not as high as the values observed in the other traits.

**Figure 1.**
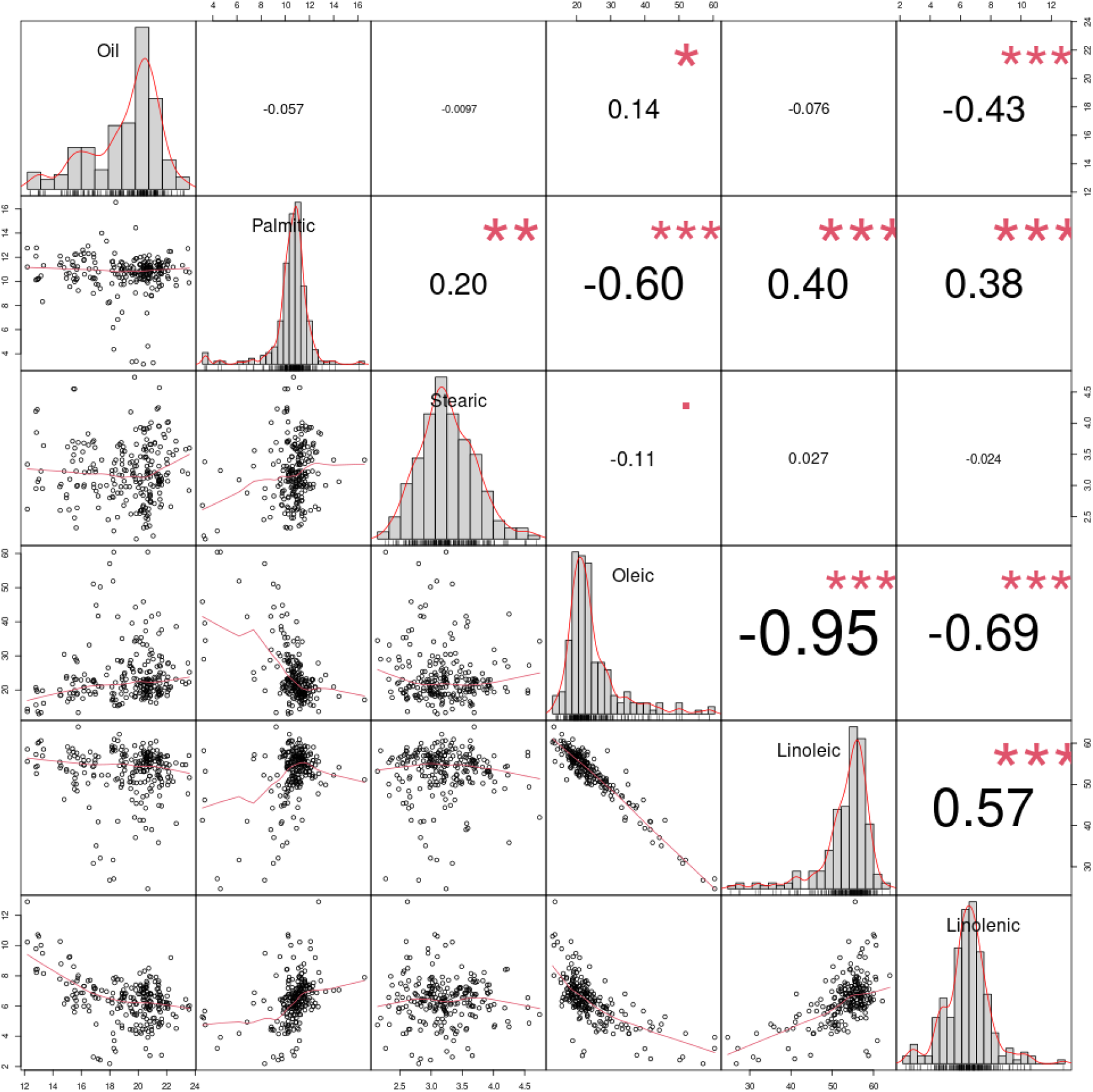
Pearson’s correlation, distribution, and phenotypic classes of the analyzed traits, total oil, oleic, palmitic, stearic, linoleic, and linolenic acids.

### Genotypic data

We used different filtering parameters to study the two genotyping approaches in the soybean oil composition. The results indicated considerable variation in SNP marker density depending on the filtering parameters selected. The total number of SNPs found in the original data by the GBS was 101304 SNPs, while for the DNA array was 6000 SNPs. The GBS method observed the largest amplitude, with an extremely low number of 177 SNPs for the 5MAF_10Het_10MD filter and a maximum of 38838 SNPS for the 5MAF_20Het_45MD filter. The number of SNPs retained after filtering by the DNA array remained more constant, with values varying from 5,403 to 6,000 SNPs depending on the filtering performed. Considering the two genotyping methodologies, the lowest total SNPs observed were with the 10% heterozygotes filter (**Table 3**). This filter is so stringent that the number of SNPs on the GBS platform is smaller than that on the DNA array platform; this is the only parameter that induces this scenario (**Figure 2**). It is essential to highlight that soybean, even though it is an autogamous plant with a smaller number of heterozygous loci, still suffers an impact on the number of SNPs with filtering for heterozygote rates, with a difference of 6648 SNPs using 10 or 20% maximum heterozygotes with GBS approach.

**Table 3.** The marker density of SNPs was observed in the genotypic data of the DNA array and GBS methodologies according to each filtering parameter.

| Filter | SNP_GBS | SNP_6K |
| --- | --- | --- |
| Original | 101304 | 6000 |
| 1MAF_10Het_10MD | 930 | 5547 |
| 1MAF_10Het_20MD | 5174 | 5573 |
| 1MAF_20Het_10MD | 1251 | 5729 |
| 1MAF_20Het_20MD | 6345 | 5758 |
| 5MAF_10Het_10MD | 177 | 5403 |
| 5MAF_10Het_20MD | 2825 | 5426 |
| 5MAF_20Het_10MD | 492 | 5585 |
| 5MAF_20Het_20MD | 3988 | 5611 |
| 5MAF_20Het_30MD | 14795 | 5617 |
| 5MAF_20Het_45MD | 38838 | 5618 |

**Figure 2.**
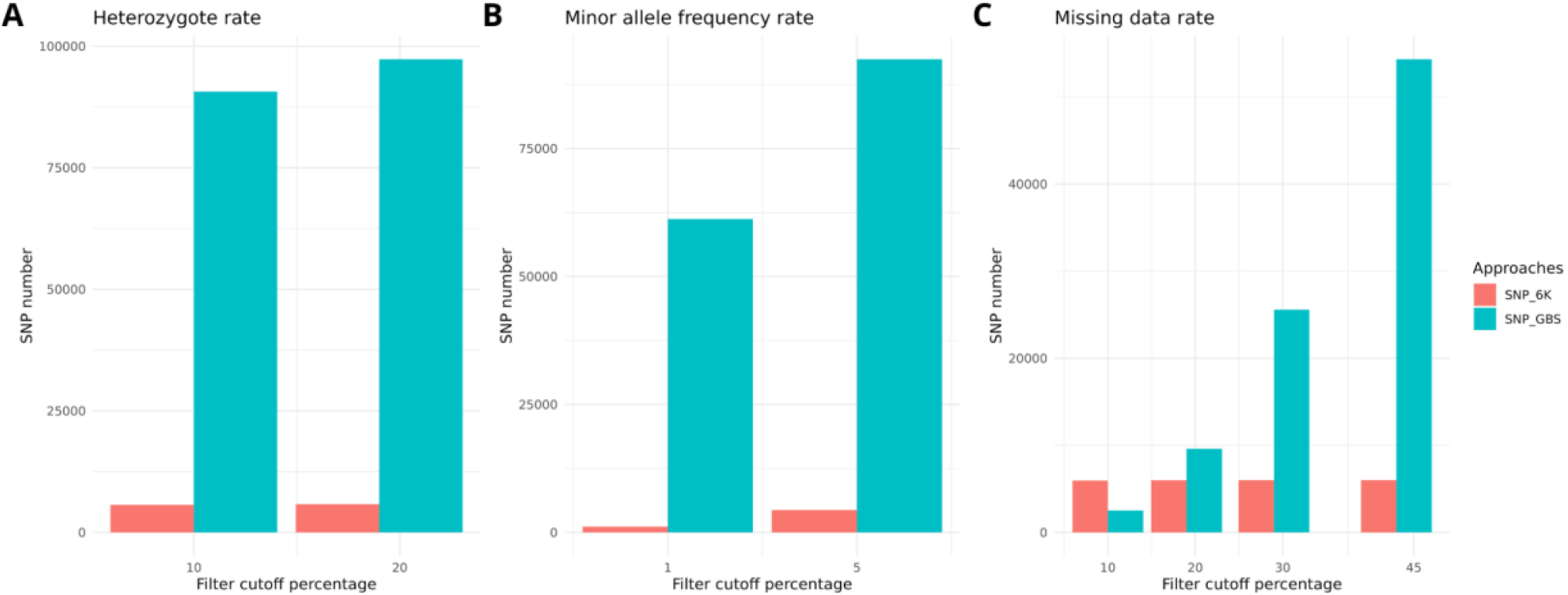
The number of SNPs in the original data before filtering and imputation, with different cutoff percentages for heterozygote rate (A), minor allele frequency rate (B), and missing data rate (C).

To analyze the population structure by the different genotyping approaches, we made a Principal Component Analysis (PCA) clustering (**Figure 3**) and a heatmap graph from the original kinship matrix (Without filters) (**Figure 4**). The kinship matrices of both genotyping approaches separated two groups. However, we observed a more precise separation of the two subpopulations in the DNA array (**Figure 4**). Furthermore, the two genotyping approaches produced different percentages of the total variance explained by the first two Principal Components (PCs); the first two of the DNA array explained 40.22% of the total variance, while the first two GBS PCs explained a total of 34.58% of the total variance (**Figure 3**). Besides, the DNA array platform divided the plant and cultivar introduction groups better than the GBS platform; there is no clear group clustering in the latter.

**Figure 3.**
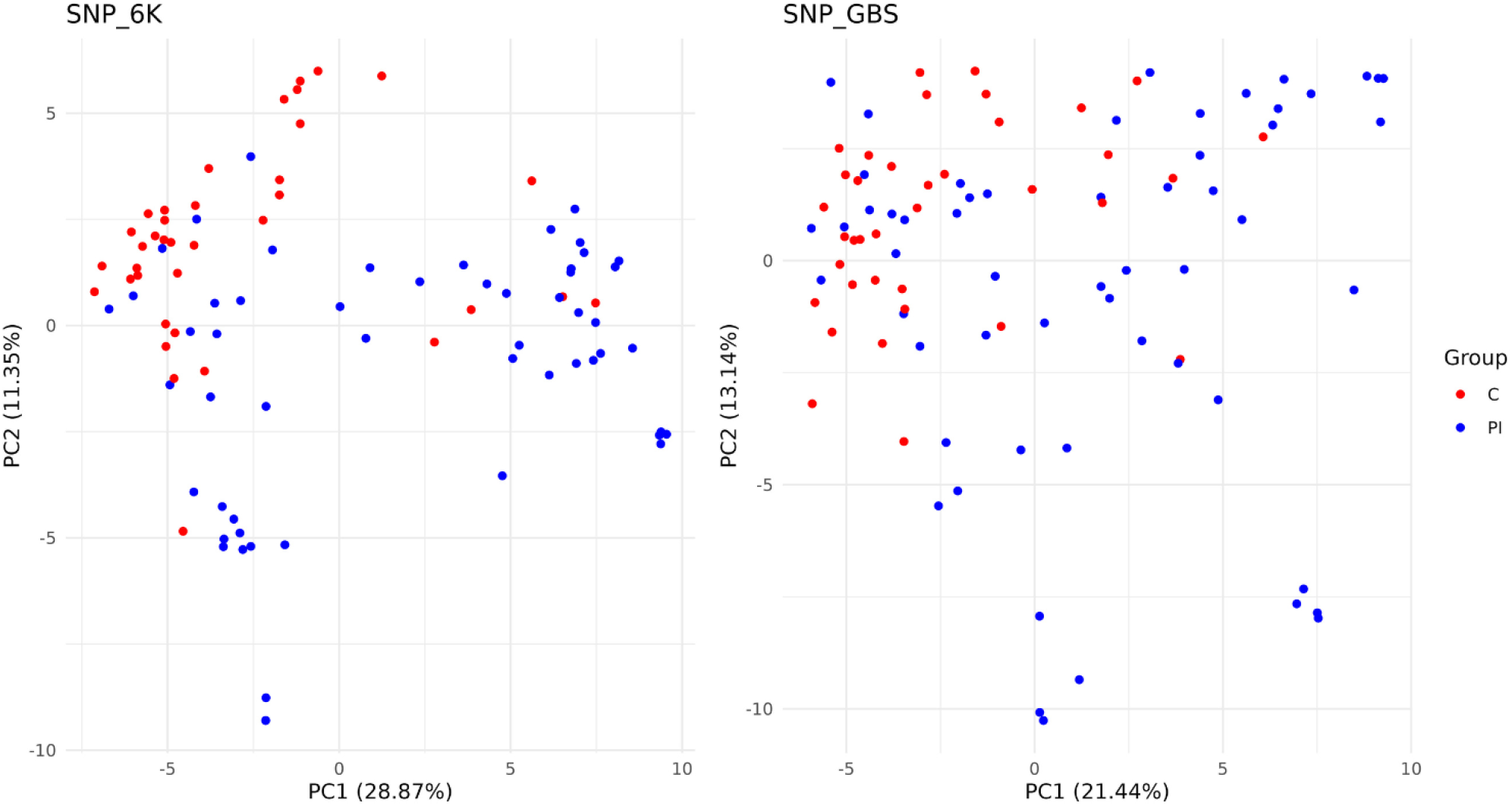
Principal Component Analysis (PCA) using kinship matrices obtained by GBS and the DNA array methodology from the original marker matrix without filters.

**Figure 3.**
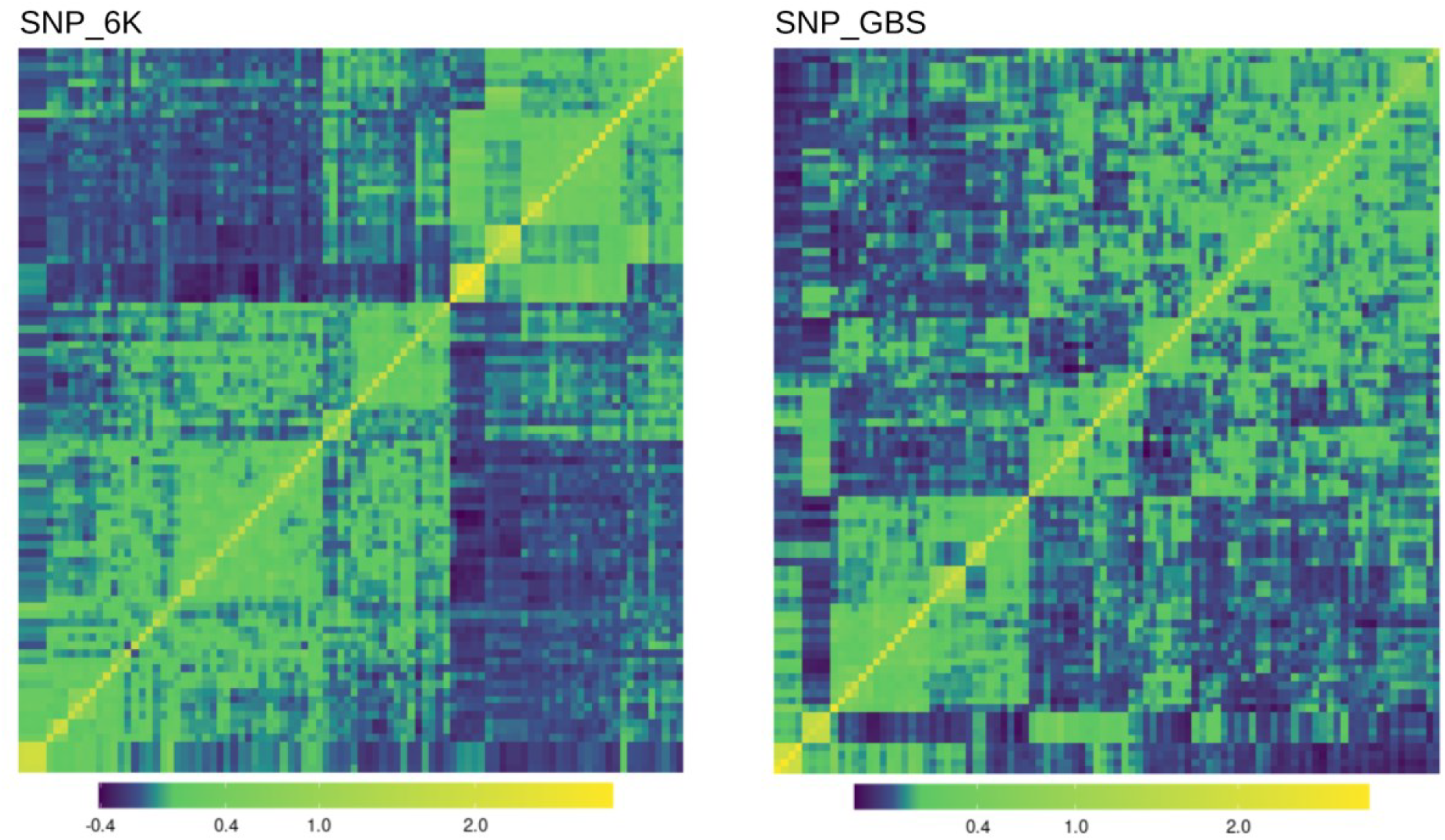
Kinship matrices using original markers obtained by GBS and the DNA array methodology from the original marker matrix without filters.

We plotted the marker density of all genomic regions using the original matrices to assess the difference in marker distribution across the genome in different genotyping platforms. The distribution of markers was different between the two genotyping approaches (**Figure 5**); the DNA array platform was shown to be denser in the telomere regions and had many gaps in the central regions. On the other hand, the GBS platform was shown to be more uniform in the distribution of its markers and not concentrated in the centromeric or telomeric regions. Furthermore, the marker-rich genomic regions of the DNA array platform have more than 19 markers, while the marker-rich regions of the GBS platform have more than 713 markers.

**Figure 5.**
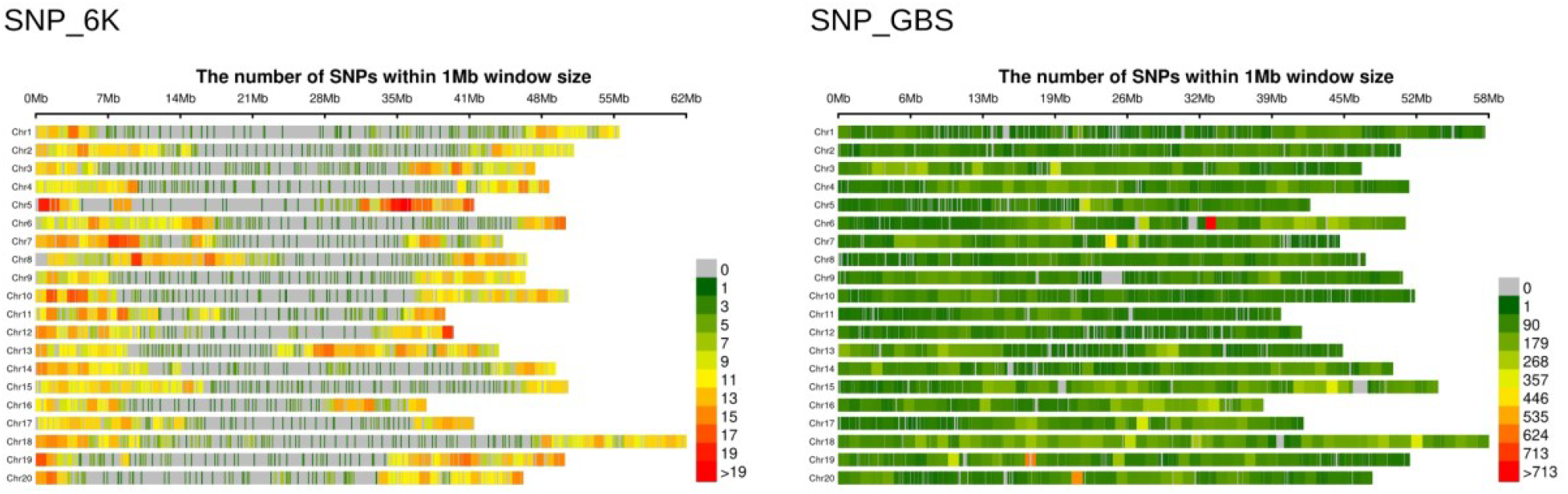
Density of SNPs markers in the original matrix of the DNA array (6000 markers) and GBS (101304).

### Prediction accuracies

Prediction accuracies for the original marker matrix in both genotyping platforms and the other filters were performed. In contrast, the predictions of all filters using “10MD” produced the lowest accuracies with GBS; this was the only one that reacted to the different filters (**Figure 5**). Furthermore, a slight difference in accuracy resulted from the heterozygotes rate only for the trait “Stearic acid”, this trait using the “1MAF_10Het_10MD” filter produced a predictive accuracy of 0.03, while the “1MAF_20Het_10MD” filter produced an accuracy of 0.17 (GBS platform and BayesB model). Minor differences were also observed for the MAF filter with the traits linolenic acid and total oil, as the former obtained a predictive accuracy of 0.41 with the “1MAF_10Het_10MD” filter, and its accuracy decreased to 0.33 when using the “5MAF_10Het_10MD” filter. Considering the average value between all filters, total oil showed the highest predictive accuracies, with mean values and standard deviations of 0.68±0.02 and 0.68±0.05 using GBS and DNA array, respectively. The trait with the lowest accuracy differed between the genotyping methods; for GBS, the trait was the palmitic acid with an accuracy value of 0.11±0.11, while for the DNA array was stearic acid with an accuracy value of 0.28±0.04. The DNA array performed better for oleic acid (accuracy value of 0.59±0.02), while linolenic and linoleic acids showed similar results in both platforms. Regarding the prediction models, the accuracies of both genotyping platforms showed similar patterns between the BayesB and BRR genomic selection models; the model’s results do not differ with the use of different filters, not even with other traits.

**Figure 5.**
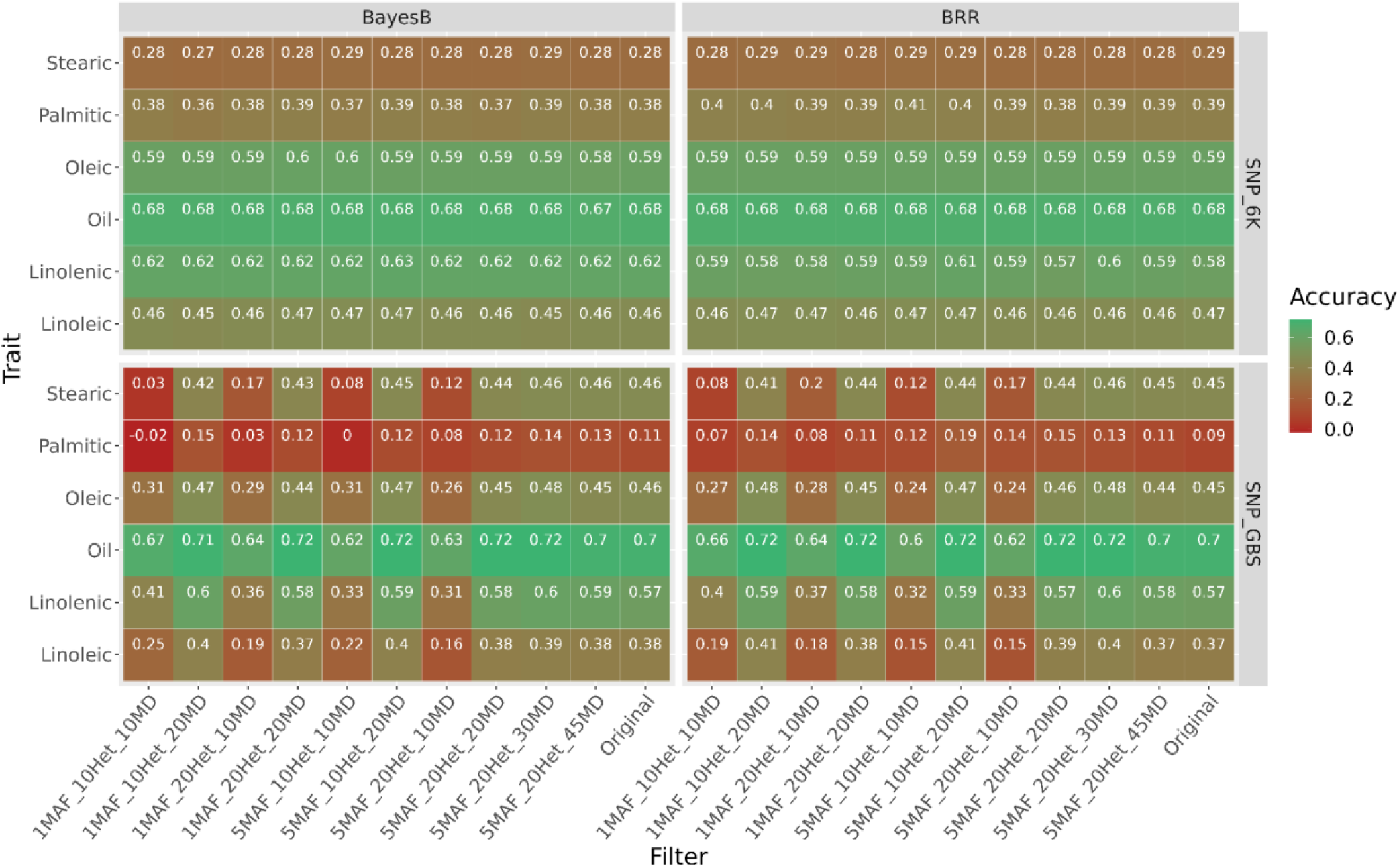
Prediction accuracies for DNA array and GBS methodology, using the BayesB and BRR genomic selection models, under different quality control parameters.

Of all the parameters used, the one that showed the most significant impact on the resulting number of SNPs was the missing data rate (**Figure 6**). Among the different rates, when the maximum 10% missing data filter was applied, the DNA array platform yielded better outcomes than GBS. From a 20% missing data rate onwards, the GBS platform outperformed the DNA array platform in predicting stearic acid and total oil traits.

**Figure 6.**
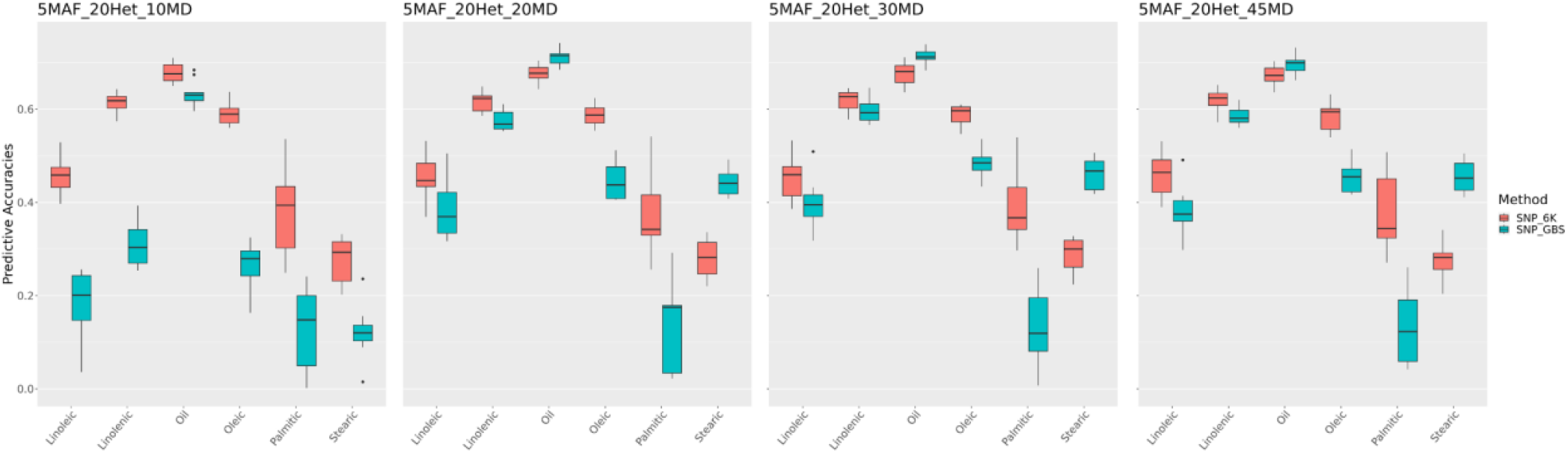
Prediction accuracy for all traits using DNA array and GBS approaches, via BayesB model, under all missing data rates, 5% of minor allele frequency and 20% of heterozygote rate.

## Discussion

Our study aimed to compare two main genotyping platforms for soybean oil composition in genomic selection. To complement the discussion on these approaches, we also used different models of genomic selection and different parameters in quality control, as the choices of these are also part of the genomic selection pipeline and may vary the predictive accuracies for each trait (ELBASYONI et al., 2018; KALER et al., 2022). The results showed that the traits predictive accuracies varied between the different genotyping platforms. Although the DNA array has important and well-studied polymorphisms for soybeans, it also has ascertainment bias since the population used for constructing the DNA array may not have the diversity comprised in the population under study. This would explain why, in our research, the GBS platform obtained better results than the DNA array platform for stearic acid; however, for palmitic and oleic acid, the accuracies were higher using the DNA array platform. In these last two traits, the diversity of the causal alleles was probably well represented in the DNA array.

In the population used, the content of oleic acid (18:1) is statistically correlated with the content of linoleic acid (18:2) and linolenic acid (18:3), specifically with a phenotypic correlation of -0.94 and -0.68. This is an expected result since oleic acid is a direct precursor of the other two acids, and some studies have already shown that when two Fatty Acid Desaturases (FADs) were knocked out, it was possible to obtain a soybean oil with up to 80% oleic acid (HAUN et al., 2014). As a result, when the oleic acid content is increased, the linoleic and linolenic acid content automatically decreases. Another high correlation found was between linolenic acid and total oil content, with a value of -0.46. Decreasing linolenic acid in soybeans disrupts the fatty acid metabolic pathway by reducing the total oil content of soybean seeds (ISLAM et al., 2019). Oleic acid (18:1) correlates -0.57 with palmitic acid (18:0) in our studied population. An experiment carried out by Zhou et al. (2021) showed that a mutant gene of the Acyl-ACP thioesterase family (FATs) resulted in a decrease in palmitic acid and an increase in oleic acid.

For soybeans, the choice of one approach or the other could be defined by the cost per sample, as the results were similar for most traits. According to studies by Priolli et al. (2019), in the same population as our study, the decay of soybean linkage disequilibrium is slow, so the gene blocks are large. Even using a population with two main groups of origin (**Figure 2**), one from Asia and the other from North and South America, it was still possible to predict the population total oil content with a predictive accuracy of 62% when using only 177 markers (filter 5MAF_10Het_10MD). The results corroborate the large gene blocks found in the linkage disequilibrium decay analysis of Priolli et al. (2019) and the only associated SNP they found for this trait. The nature of the soybean crossing can also explain this fact, as it is an inbred crop. **Figure 3** shows how the GBS platform markers can be well distributed throughout the genome, but this characteristic does not confer superior accuracy for most traits. For example, Yu et al. (2023) obtained superior accuracies for all traits using GBS and maize, an outbreed crop. For an outbred population, the filters would probably generate more diverse results in accuracy due to the high number of recombination points, more considerable number of heterozygous loci, and shorter gene blocks.

The filters that obtained the lowest values in predictive accuracies for this soybean population were all those that contained the maximum of 10% of missing data allowed. However, these differences mainly affected the GBS platform because the accuracies of the DNA array platform remained the same with the different applied filters. This can also be seen in the results obtained by Elbasyoni al. (2018), in which all accuracies by the GBS platform with a maximum missing data value of 50% outperformed all predictions made with a maximum of 10% MD filtering in Winter Wheat. On the other hand, the different filtering and genotyping approaches did not result in contrasting predictive accuracies when using different predictive models. We expected that the BayesB model would result in higher predictive accuracy in our case because the model considers many loci with no effect on the trait. There are higher effects alleles, and as mentioned before, oil composition traits have alleles with higher effects. However, like Moraes et al., 2018 and Silva, Xavier, Faria (2021), different predictive models resulted in similar accuracies. Furthermore, the small population used may have reduced statistical power, consequently increasing difficulty in differentiating the predictive accuracies for each model.

Although our work has a small population for genomic prediction, it represents an essential study on the genotyping platforms currently used in traits that are costly to phenotype and of great importance for soybean breeding since soybean oil is one of its main final products. Furthermore, Jarquín et al. (2014) showed that a small to moderate training set is enough to achieve high accuracy in soybean genomic prediction, stabilizing around a training population size slightly bigger than 100 for grain yield. Moreover, considering that traits are highly correlated (**Figure 1**), that genotyping platforms performed differently between traits, and that quality control also impacted accuracy, our work has a vital role in streamlining and directing the work of breeders. The GBS platform or the DNA array may be more suitable depending on the target oil characteristics a breeding program searches for. For example, if the objective is to produce a healthier oil with lower stearic acid content (LEAMY et al., 2017), the GBS platform was able to deliver higher accuracy than *BarcSoySNP6k*.

Furthermore, stearic acid only has a significant correlation of 20% with palmitic acid (another saturated acid), so indirect selection using other traits is impossible for this acid. Conversely, the DNA array has higher accuracies for most traits. It has significant stability and is not prone to quality control error, making it an excellent choice for general trait improvement. Total oil, a less costly trait than traits derived from the oil’s chromatographic profile, has a significant correlation of 43% with linolenic acid. This high correlation could indicate potential use in indirect selection, especially in preliminary phases of line evaluation. Additionally, in the case of the DNA array platform, no filtering and only imputation would be choices that would lead to solid results. In contrast, the “5MAF_20Het_30MD” filter and imputation would be the most suitable choice for the GBS platform, as it has obtained the highest accuracy.

In conclusion, the present analysis confirmed the utility of the DNA array and GBS platform for prediction analysis and its direct applicability for soybean improvement. By using favorable SNP markers, soybean breeders can rapidly improve oil traits in soybean programs.

## Supplementary material

**Supplementary 1.** Published papers with genomic prediction in soybeans, their genotyping approaches, and quality control parameters.

| Article | Genotyping method | Filters | Year | DOI |
| --- | --- | --- | --- | --- |
| Accuracy of genomic prediction for seed oil concentration in high-oleic soybean populations using a low-density marker panel | * | Missing data<30%;<br>MAF <=10% | 2021 | 10.1002/csc2.20607 |
| Cold stress tolerance of soybeans during flowering: QTL mapping and efficient selection strategies under controlled conditions | GBS | SNP call rate > 85%;<br>missing values MAF < 0.375; Heterozygosity > 10% | 2019 | 10.1111/pbr.12734 |
| Context-Specific Genomic Selection Strategies Outperform Phenotypic Selection for Soybean Quantitative Traits in the Progeny Row Stage | DNA array | Duplicate markers;<br>Identify likely genotyping errors by LOD scores | 2019 | 10.2135/cropsci2018.03.0197 |
| Genomic prediction using training population design in interspecific soybean populations | DNA array | SNPs call rate < 90% | 2021 | 10.1186/1746-4811-9-44 |
| Genomic Selection for Yield and Seed Composition Traits Within an Applied Soybean Breeding Program | DNA array | Monomorphic SNPs;<br>heterozygotes > 10%;<br>Missing data >80%;<br>MAF <0.05 | 2019 | 10.1534/g3.118.200917 |
| Genomic Selection for Yield and Seed Protein Content in Soybean: A Study of Breeding Program Data and Assessment of Prediction Accuracy | DNA array | SNPs call rate >20%;<br>Heterozygosity > 15%;<br>MAF > 0.05; | 2017 | 10.2135/cropsci2016.06.0496 |
| Genomic selection in soybean: accuracy and time gain about phenotypic selection | DNA array | 0.01 < MAF > 0.99;<br>SNPs call rate > 10%<br>missing data removed | 2018 | 10.1186/1297-9686-46-32 |
| Genomic selection performs as effectively as phenotypic selection for increasing seed yield in soybean | GBS | Percentage of missing values <= 80%;<br>MAF<0.01 | 2022 | 10.1002/tpg2.20285 |
| Genotyping by sequencing for genomic prediction in a soybean breeding population | GBS | MAF 0.01 ;Duplicate sites; | 2014 | 10.1186/1471-2164-15-740 |
| Improvement of key agronomical traits in soybean through genomic prediction of superior crosses | GBS | Missing data >80%;<br>Minor Allele Count <4;<br>Heterozygotes >5.4%; | 2020 | 10.1002/csc2.20583 |
| Optimizing genomic selection in soybean: An important improvement in agricultural genomics | GBS | MAF 0.05 | 2022 | 10.1016/j.heliyon.2022.e11873 |
| Selection of soybean elite cultivars based on phenotypic and genomic traits related to lodging tolerance | DNA array | MAF >0.05; SNPs call rate <0.25 | 2017 | 10.1111/pbr.12495 |
| Testing methods and statistical models of genomic prediction for quantitative disease resistance to Phytophthora sojae in soybean [Glycine max (L.) Merr] germplasm collections | DNA array | Monomorphic;<br>MAF<0.05 | 2020 | 10.1007/s00122-020-03679-w |

\* No information

